# Miswired enhancer logic drives translocation positive rhabdomyosarcoma

**DOI:** 10.1101/757369

**Authors:** Berkley E. Gryder, Marco Wachtel, Kenneth Chang, Osama El Demerdash, Nicholas G. Aboreden, Wardah Mohammed, Winston Ewert, Silvia Pomella, Rossella Rota, Jun S. Wei, Young Song, Beat Schaefer, Christopher R. Vakoc, Javed Khan

**Affiliations:** Genetics Branch, National Cancer Institute, NIH, Bethesda, MD 20892, USA; University Children’s Hospital, Zurich, Switzerland; Cold Spring Harbor Laboratory, 1 Bungtown Road, Cold Spring Harbor, NY 11724, USA; Biologic Institute, Redmond, Washington, USA; Department of Oncohematology, Ospedale Pediatrico Bambino Gesu’ Research Institute, IRCCS, Rome, Italy

**Keywords:** 3D Chromatin, Enhancer Logic, Childhood Cancer, Transcription Factors

## Abstract

Core regularity transcription factors (CR TFs) define cell identity and lineage through an exquisitely precise and logical order during embryogenesis and development. These CR TFs regulate one another in three-dimensional space via distal enhancers that serve as logic gates embedded in their TF recognition sequences. Aberrant chromatin organization resulting in miswired circuitry of enhancer logic is a newly recognized feature in many cancers. Here, we report that *PAX3-FOXO1* expression is driven by a translocated *FOXO1* distal super enhancer (SE). Using 4C-seq, a technique detecting all genomic regions that interact with the translocated *FOXO1* SE, we demonstrate its physical interaction with the *PAX3* promotor only in the presence of the oncogenic translocation. Furthermore, RNA-seq and ChIP-seq in tumors bearing rare PAX translocations implicate enhancer miswiring is a pervasive feature across all FP-RMS tumors. HiChIP of enhancer mark H3K27ac showed extended connectivity between the distal *FOXO1* SE and additional intra-domain enhancers and the *PAX3* promoter. We show by CRISPR-paired-ChIP-Rx that *PAX3-FOXO1* transcription is diminished when this network of enhancers is selectively ablated. Therefore, our data reveal a mechanism of a translocated hijacked enhancer which disrupts the normal CR TF logic during skeletal muscle development (PAX3 to MYOD to MYOG), replacing it with an infinite loop logic that locks rhabdomyosarcoma cells in an undifferentiated proliferating stage.

## Introduction

Control of the expression of the core regulatory transcription factors (CR TFs) that guide developmental decision making are directed by logical enhancer elements^1–3^. These genomic elements, when heavily activated, become super enhancers (SEs) with unusually large deposits of active histone marks, chromatin regulators and transcriptional coactivators^4^. Chromosomal rearrangements allowing SEs to drive oncogene expression is an emerging mechanism in tumor biology^5,6^. Alveolar (fusion-positive) rhabdomyosarcoma (FP-RMS), an aggressive skeletal muscle cancer of childhood, often possesses chromosomal translocations, involving commonly *PAX3* and *FOXO1* genes, rarely *PAX7*-*FOXO1*, and in exceptional cases *PAX3*-*INO80D* and *PAX3*-*NCOA1* fusions^7^. Disruption of CR TF transcription is effectual as FP-RMS treatment^8,9^. During normal skeletal muscle development, PAX3 initiates specification of the muscle lineage and is shut off during myogenic differentiation. Consequently master regulators MYOD and finally MYOG promote muscle progenitor cells to exit cell division and complete muscle differentiation^10^. However, although FP-RMS cells express these master regulators needed to trigger muscle differentiation program, they are halted in an early myoblastic and thus more proliferative state and not able to complete cell differentiation. Fusion gene product resulted from the translocation has been thought to be responsible of the inability of differentiation for the FP-RMS. However, the mechanism of how the oncogenic fusions lock FP-RMS cells in their myoblast state has not been fully understood. In this study, we test the hypothesis that the chromosomal translocation event resulted in novel enhancer/promoter interactions to maintain robust expression of the oncogenic fusion protein in FP-RMS.

When conducting the first epigenetic landscape study of FP-RMS we uncovered a strong dependence on SEs for tumor survival, with PAX3-FOXO1 being a chief determinant of SE formation in collaboration with MYOD and MYOG, and oncogene MYCN^9^. Using chromatin conformation capture (3C, 4C-seq) and chromatin immunoprecipitation (ChIP-seq, ChIP-Rx, HiChIP) based assays, we here study a key SE 300 kb distal of *FOXO1* which is occupied by all four of these master regulators, and examine its function in FP-RMS. We propose that hijacking SEs bound by myogenic CR TFs allows for continued expression of oncogenic *PAX* fusions, thus circumventing normal myogenic enhancer logic.

## Results

### Chromosomal translocation imports the *FOXO1* super enhancer to the *PAX3* promoter

Precisely how PAX3-FOXO1 locks the cells into a myoblastic state unable to differentiate is unknown. Proper enhancer-promoter interactions are enabled by constraints in 3D chromatin folding, determined by CTCF and cohesion formed loops at topologically associated domain (TAD) boundaries^11,12^. *PAX3* is normally silenced during progression past the myoblast stage of muscle differentiation. PAX3 expression during embryogenesis is tightly controlled, and structural variation that disrupts the PAX3 TAD causes limb malformation^13^. We hypothesized that the fusion event results in novel enhancer/promoter looping events to maintain fusion protein expression independent of normal lineage control. Hi-C data^14^ indicated 3 candidate topological loops containing wild-type *FOXO1* that exist in normal cells. We found all of these were occupied by RAD21 (of the cohesion complex) and CTCF in FP-RMS RH4 cells by ChIP-seq (Fig. 1a). CTCF binding events that form loops most often have binding motif sequences that are antiparallel (and point inward toward each other)^14^. The CTCF motif orientation at the 1^st^ and 3^rd^ of these sites near *FOXO1* were found to be antiparallel with the CTCF motif near the PAX3 promoter, permissive of chromatin loop formation via extrusion after the translocation.

**Figure 1.**
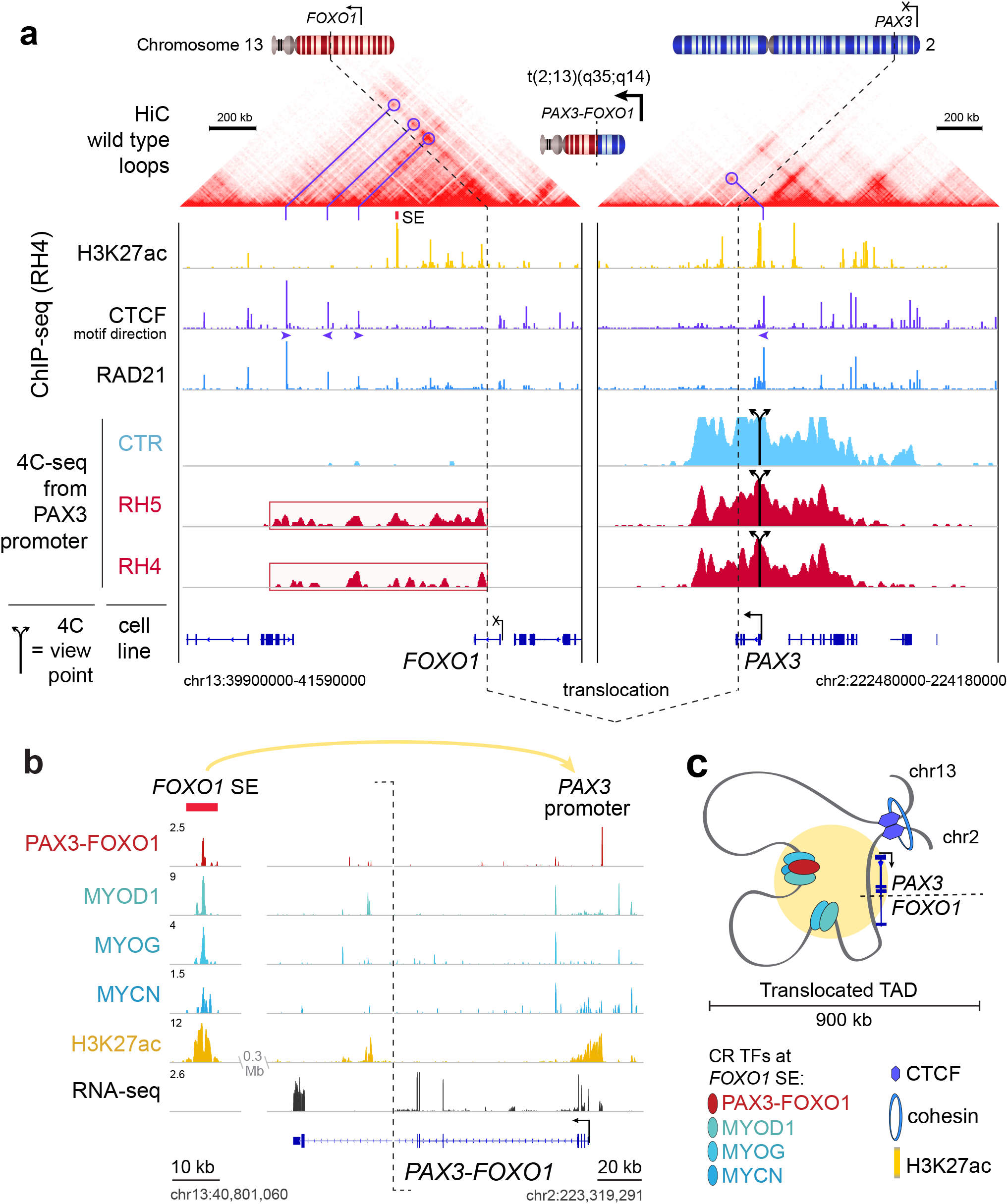
Translocation restructures insulated neighborhood surrounding PAX3-FOXO1. **a.** Wild-type loops indicated by Hi-C profile from human GM12878 cells. ChIP-seq demonstrates binding locations of H3K27ac, CTCF and RAD21 in RH4 cells. 4C-seq reveals looping between viewpoints at CTCF sites bounding FOXO1 enhancers, and the PAX3 promoter, in translocation negative (CTR) and translocation positive (RH5, RH4) cells. Viewpoints are indicated by split arrows, and translocation breakpoints are indicated by dotted lines. **b.** ChIP-seq signal for Master Transcription Factors and H3K27ac, and RNA-seq signal, in Reads Per Million (RPM), at the FOXO1 Super Enhancer (SE) and PAX3-FOXO1 fusion gene. **c.** Schematic of translocation created a new Topologically Associated Domain (TAD) bringing the *PAX3*-promoter (chr2) under the control of *FOXO1* SE and other smaller enhancers (chr13).

To identify interacting domains *cis* to the *PAX3* promoter after the translocation, we used circularized chromatin conformation capture followed by sequencing (4C-seq) from viewpoint anchors around the *PAX3* promoter and *FOXO1* genes on chromosomes 2 and 13.

Remarkably, looping was detected between the *PAX3* promoter and multiple candidate enhancers downstream of *FOXO1*, and was restricted between the intronic fusion breakpoint in *FOXO1* and the predicted topological boundary (Fig. 1a). The outermost TAD-boundary looping interaction was confirmed by Sanger sequencing of the 3C PCR product (Extended Data Fig. 1a-c). Notably, each of the 3 CTCF sites 3’ of *FOXO1* formed looping interactions with *PAX3* only in translocation-positive RH4, but not in the translocation-negative RMS cell line CTR (Extended Data Fig. 1d). A previous study of 4C-seq in FP-RMS cell lines has shown similar interactions consistent with our results^15^, but here we provide the first ChIP-seq informed functional interpretation of these interactions as CTCF bounded enhancers that contain critical CR TFs. We hypothesize these newly juxtaposed enhancer elements keep the *PAX3* promotor on the translocated allele active. These enhancers could sustain the continual *PAX3-FOXO1* expression and the oncogenic process because they harbor strong binding sites for MYOD1, MYCN and MYOG (Figure 1b-c).

### Rare PAX-fusions implicate enhancer miswiring

Besides the *PAX3-FOXO1* translocation, there are several other PAX translocation variations in FP-RMS including *PAX7-FOXO1*, *PAX3-NCOA1*, and *PAX3-INO80D* (Figure 2a). While there is protein homology between PAX3 and PAX7 (with similar DNA binding domains), NCOA1 or INO80D share hardly any protein homology with the FOXO1 transactivation domain. However, RNA-seq reveals that remarkably similar transcriptome profiles from tumors harboring these diverse oncogenic fusions^7^. Additionally, SEs (found in RH4 cells to be bound by FP-RMS specific CR TFs) exist near all known translocation partner genes (Figure 2b). This led us to hypothesize that enhancer miswiring as a result of chromosomal translocations may be the common theme among all PAX-fusions tumors.

**Figure 2.**
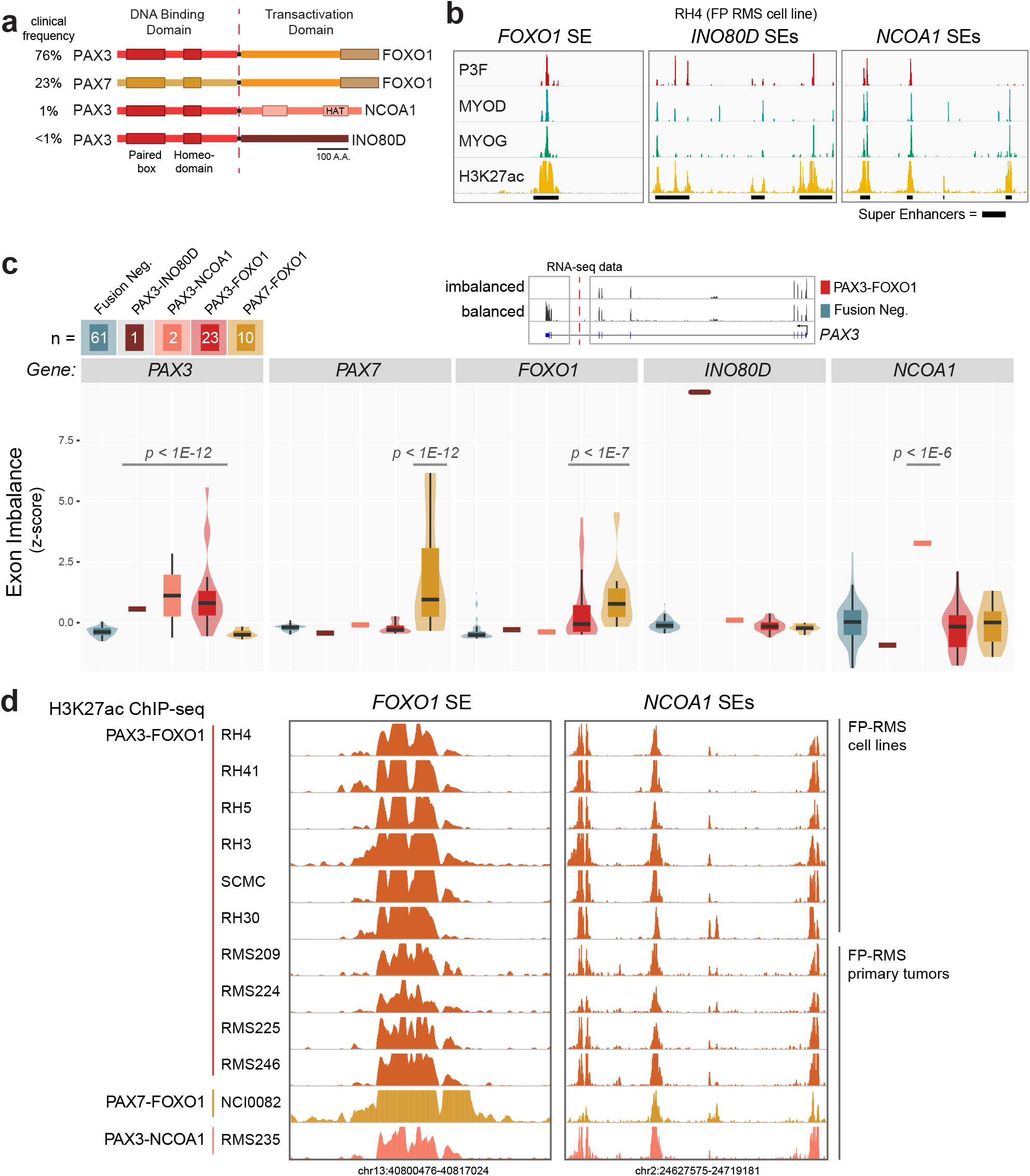
SEs and allele specific expression at rare PAX3 translocation partners. **a.** PAX fusions and their clinical frequency in FP-RMS patient tumors. **b.** SEs in RH4 (PAX3-FOXO1 bearing cells) include not only those near *FOXO1*, but also *INO80D* and *NCOA1*. **c.** Exonic imbalance measure in RNA-seq data from primary tumors and cell lines of FN-RMS (n = 61), FP-RMS with PAX3-INO80D fusion (n = 1), PAX3-NCOA1 fusion (n = 2), PAX3-FOXO1 (n = 23) and PAX7-FOXO1 (n = 10). P-values were calculated using a two-tailed t-test with Welch’s correction. **d.** H3K27ac ChIP-seq in FP-RMS cell lines (n = 6) and primary tumors (n = 6) at the *FOXO1* SE and the *NCOA1* SEs.

In FP-RMS with a *PAX3-FOXO1* translocation, if only the promoter of *PAX3* determines the expression of the *PAX3* gene on the wild-type allele and *PAX3-FOXO1* fusion gene on the translocated allele, the expression of *PAX3* exons will be even. The expression of *PAX3* exons of will be uneven if *PAX3-FOXO1* is regulated by the abnormal juxtaposition of the *FOXO1* SE, since the last exons of *PAX3* are not influenced by the *FOXO1* SE (both from the remaining wildtype *PAX3* and the reciprocal *FOXO1-PAX3* translocated allele). Therefore, we examined exon level expression of the genes involved in translocation using RNA-seq data from FP-RMS patient tumors. The RNA-seq data showed that exons before the translocation (3-prime or N-terminal) are always expressed significantly higher than those beyond the translocation breakpoint (5-prime or C-terminal) (Figure 2c). For example, exon-level expression in tumors with *PAX3-FOXO1* revealed that the final two post breakpoint exons of *PAX3* were greatly under expressed only in the tumors with *PAX3* fusion genes, indicating that only the exons involved in the fusion event is expressed due to activation by the *FOXO1* SE. On the other hand, exon usage of a gene is balanced in RMS patients lacking the translocation, e.g. *PAX3* in fusion negative RMS (Figure 2c). Importantly, inferring allele selective expression via RNA-seq allows interrogation of extremely rare PAX fusions, *PAX3-INO80D* and *PAX3-NCOA1*. All fusion gene partners showed exonic imbalance resulting from favored expression of the translocated exons (Fig 2c, Extended Data Fig 2).

ChIP-seq data in FP-RMS cell lines and patients allowed us to discover recurrent SEs surrounding not only *FOXO1*, but also rare partner *NCOA1* (Figure 2d). Although SEs represent only ~4% of enhancers, their unique and consistent presence argues that these diverse fusions may uniformly rewire SEs which can be active in the epigenomic state of all FP-RMS patients. Here we report the first epigenomic data generated for a PAX3-NCOA1 patient, and we found this rare fusion epigenetically phenocopies tumors with PAX3-FOXO1 or PAX7-FOXO1 (Figure 2d). When juxtaposed, these recurrent SEs are key elements driving the expression of fusions genes.

### CRISPR reveals essentiality of cis-regulatory elements regulating PAX3-FOXO1

To build on the evidence from 4C, we set out to gain a more complete dataset confirming the interactions between the enhancer network controlling *PAX3-FOXO1*. Thus, we used HiChIP against H3K27ac to capture protein-directed interaction frequency between acetylated chromatin sites at enhancers and promoters^16^. The results identified that the *FOXO1* super enhancer is connected not only to the *PAX3* promoter, but also to three smaller intergenic and intronic enhancer elements near or within *FOXO1* (Figure 3a-b).

**Figure 3.**
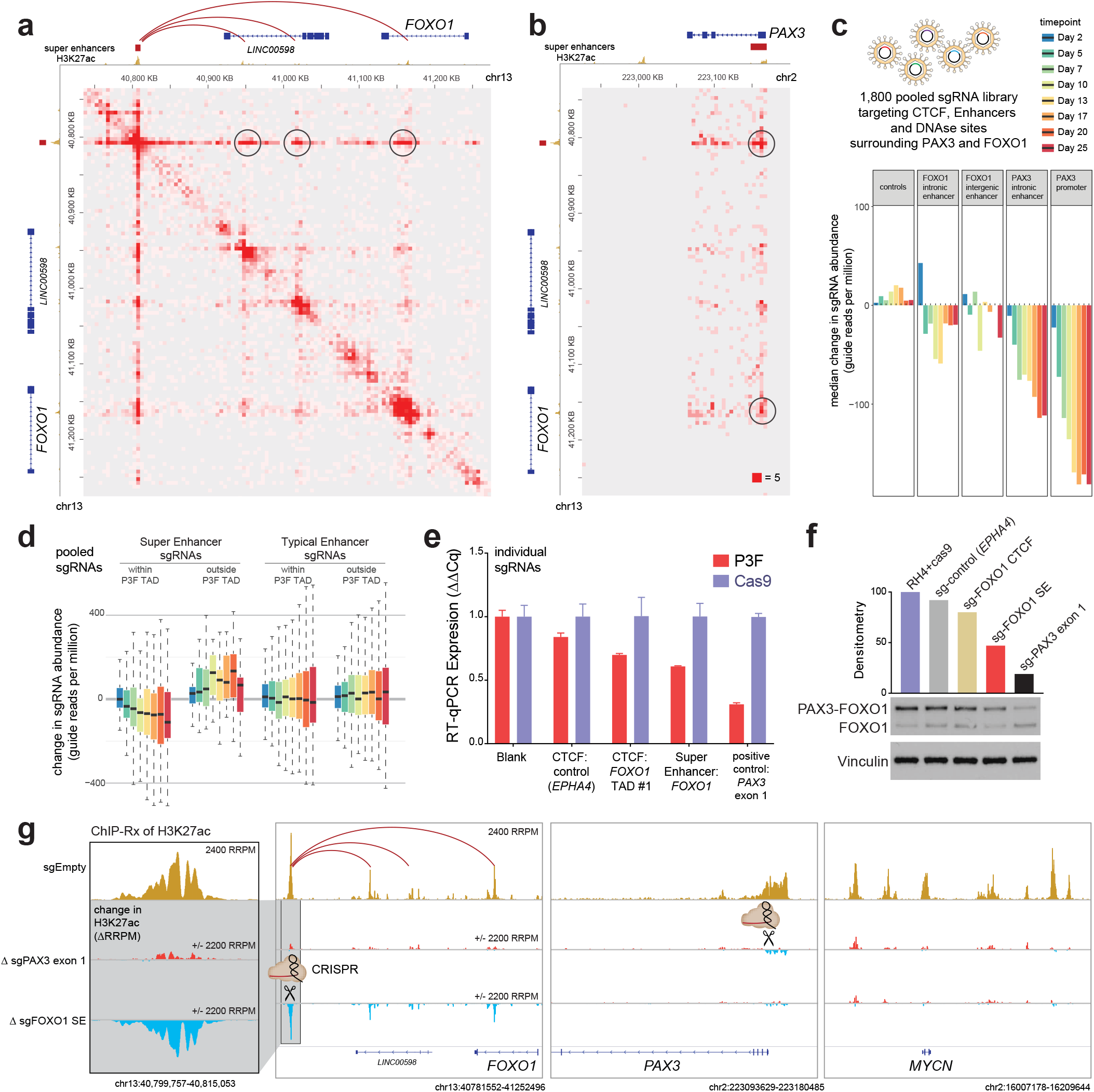
CRISPR functional mapping of non-coding elements controlling PAX3-FOXO1. **a.** H3K27ac HiChIP reveals structure of FOXO1 SE interactions with smaller intra-TAD enhancer elements. **b.** Interaction by H3K27ac HiChIP between PAX3 promoter and FOXO1 SE and intronic enhancer element. **c.** Pooled sgRNA CRISPR screening tiling against *cis*-regulatory genomic elements surrounding *PAX3* and *FOXO1* defined their degree of essentiality. RH4 cells expressing Cas9 were sampled by counting sgRNA abundance using sequencing at the indicated time intervals. **d.** Change in sgRNA abundance of pooled CRISPR shows intra-TAD super enhancers are more critical for RH4 cell survival than typical enhancers, or SEs outside TAD boundaries. **e.** Individual sgRNA impact on PAX3-FOXO1 gene expression after 1 day of transduction in FP-RMS cells RH4 expressing cas9. **f.** PAX3-FOXO1 protein levels are reduced by individual sgRNAs targeting key *cis*-regulatory elements, especially those targeting the *FOXO1* SE and the first exon of *PAX3*. **g.** ChIP-seq with reference exogenous spike-in (ChIP-Rx) for H3K27ac was employed to interrogate the chromatin impact of the sgRNA targeting the *FOXO1* SE. The top track is control ChIP-Rx (sgEmpty), the second track is showing the change (delta RRPM) upon CRISPR of the 1^st^ exon of *PAX3*, and the third track shows the change in H3K27ac upon CRISPR of the *FOXO1* SE at 24 hours post sgRNA transduction. All experiments were performed in RH4 cancer cells.

We next measured the contribution of these enhancer elements to the overall survival of FP-RMS cells. We designed a library of sgRNAs against each enhancer or promoter constituent, each DNase-hyper sensitive site and each CTCF peak as defined by genome-wide profiles in RH4 cells. We introduced them in a pooled fashion by viral infection into RH4 cells expressing Cas9. The abundance of each sgRNA in the population was then quantified over time using next-generation sequencing (at day 2, 5, 7, 10, 13, 17, 20 and 25). sgRNA’s that target the *PAX3* promoter had the strongest impact on RH4 cell viability, as inferred from the largest reduction in guide representation over time (Figure 3c). Among CTCF sites, two candidate anchor sites (*FOXO1*-distal sites #2 and #3) had no negative influence, while the outermost CTCF sites (*FOXO1*-TAD boundary site #1 and *PAX3*-TAD boundary) were both reduced by negative selection (Extended Data Figure 3). Among TF-bound and H3K27ac decorated enhancers, sgRNA’s targeting SEs within the TAD (the translocation-induced insulated neighborhood containing *PAX3*-*FOXO1*) were more effective than sgRNA’s targeting SEs outside this neighborhood, and also more than typical enhancers (Figure 3d).

Individual sgRNA’s were next used to study the impact on *PAX3*-*FOXO1* transcription. We confirmed our hypothesis that disruption of the *FOXO1* SE reduced PAX3-FOXO1 at the transcript level and protein level at 24 hours after sgRNA infection (Figure 3e-f). To attribute the effect of this sgRNA to direct impairment of the enhancer, we assayed H3K27ac changes by ChIP-Rx (spike in reference normalized ChIP-seq)^17^. The results revealed that the *FOXO1* SE was not only depleted of H3K27ac, but that the associated enhancer interaction network (as HiChIP identified) was also drastically reduced of acetylation at the sites interacting with *FOXO1* SE (Figure 3g). Conversely, this enhancer network was not impaired by a sgRNA targeting the first exon of *PAX3*, except for slight reduction in acetylation levels at the *PAX3* promoter (Figure 3g). These data demonstrated that the *FOXO1* SE is essential to maintain the expression of *PAX3-FOXO1* oncogene in RMS.

### *FOXO1* SE is activated during a key step in myogenesis

To examine if the activity of the *FOXO1* SE was coordinated with myogenic steps, we utilized ENCODE data mapping H3K27ac in various stages of the muscle lineage. We found that the *FOXO1* enhancer is transiently transformed into a super enhancer during myogenesis at the same time *MYOG* acquires a super enhancer (Figure 4a). *FOXO1*, *MYOG*, and *MYOD1* have more highly ranked SEs in FP-RMS as compared to FN-RMS (Figure 4b), in agreement with the notion that FP-status is more advance toward myotubes, and FN-status is more similar the earlier myoblast state. *MYOG* activation is commonly prevented by mutant-RAS signaling through MEK/ERK in FN-RMS tumors, which can be rapidly release *via* small molecule inhibitors of MEK/ERK^18^. Using this system, we asked if the *FOXO1* SE and concomitant *FOXO1* expression was induced alongside *MYOG* activation. Indeed, not only did we find *FOXO1* to be upregulated, but also observed MYOG invasion on the same SE which is recruited during the PAX3-FOXO1 translocation event in FP-RMS (Extended Data Fig. 4a).

**Figure 4.**
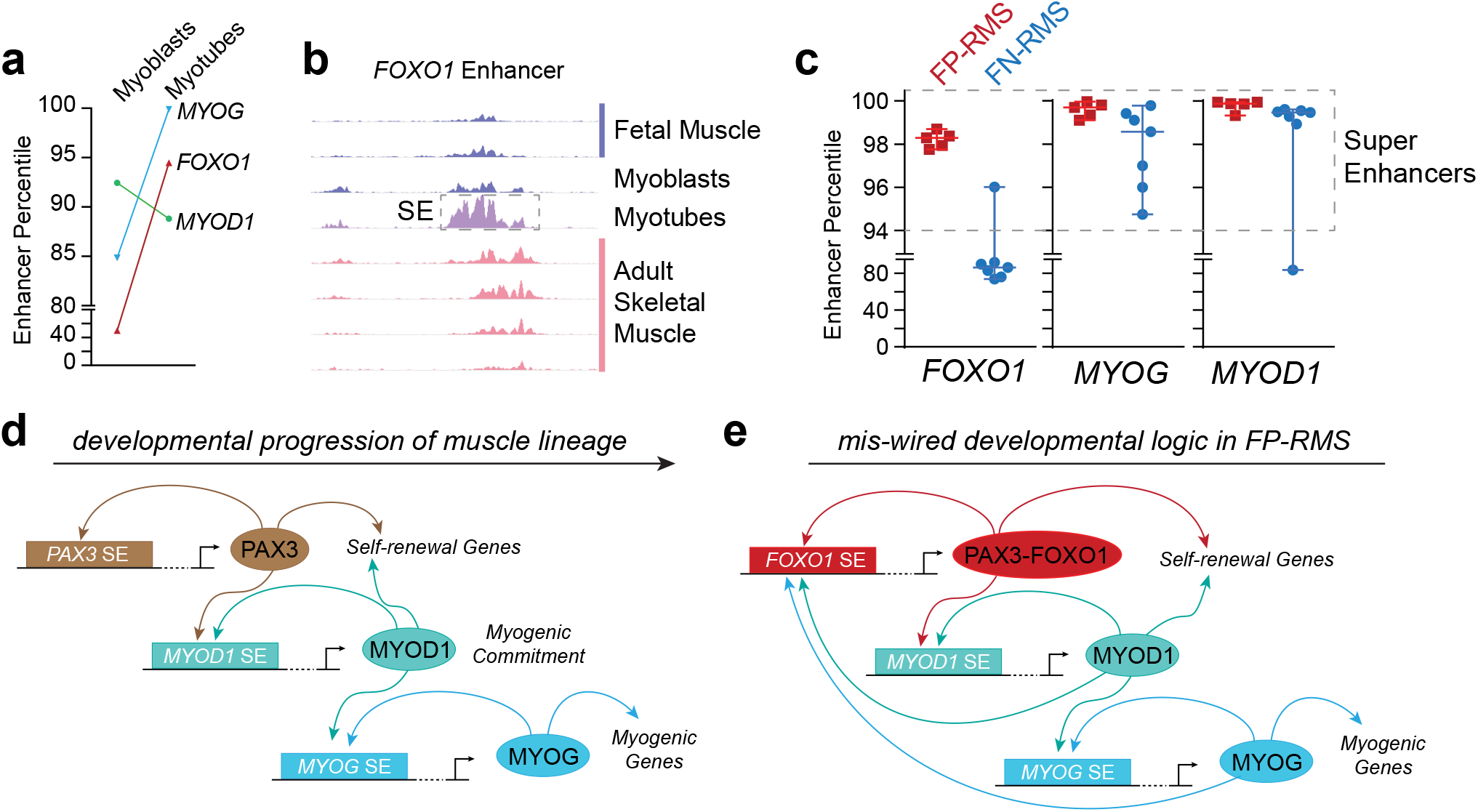
Miswired super enhancer logic able to maintain an oncogenic cell state. **a.** Increased FOXO1 and MYOG enhancer rank (by percentile) in the transition from myoblasts to myotubes. **b.** Enhancer rank of *FOXO1*, *MYOG* and *MYOD1* in RMS, as measured by rank of H3K27ac ChIP-seq bound to each enhancer. **c.** Model illustrating normal development of the muscle lineage. **d.** Miswiring of myogenic circuitry due to translocation of FOXO1 SE to PAX3 promoter, allowing MYOD and MYOG to activate PAX3-FOXO1.

## Discussion

Two factors can be selected for in rearrangement driven cancers: oncogenic biochemical function in the case of a resulting fusion protein^19^, or aberrant expression levels of a proto-oncogene (such as *MYC* or *GFI1*) via enhancer hijacking^5^. In FP-RMS, PAX-fusions are selected for both an oncogenic fusion protein product and miswired enhancer logic, effectively reprograming core regulatory TF networks.

PAX3 activates MYOD *via* binding and activating *MYOD1* SEs, but then shuts off presumably because MYOD does not work backwards to upregulate *PAX3* (MYOD ChIP-seq shows no binding in the *PAX3* promoter, unlike the *MYOG* promoter, Extended Data Fig. 4b). Lacking enhancers responsive to MYOD/MYOG, the remaining wild-type alleles of *PAX3/7* in FP-RMS tumors are silent. Our data suggests that newly juxtaposed enhancer elements initiate and continually drive *PAX3-FOXO1* expression, implicating that enhancer miswiring is at the heart of the oncogenic process in FP-RMS. When the *FOXO1* SE is translocated to regulate *PAX3*, late myogenic factors (MYOG/MYOD) work through this SE to drive an early myogenic factor (PAX3), changing a “progressive” enhancer logic into an “infinite loop” enhancer logic. Similarly, CR TF logic circuits are self-reinforcing in self-renewal and proliferative states such as embryonic stem cells^20^.

Analysis of RNA-seq for patients with non-FOXO1 partners (INO80D, NCOA1) of PAX3 reveals a remarkably similar transcriptome^7^, despite not being TFs themselves, and having no sequence homology to the activation domain (AD) of FOXO1. It has been shown that TFs can perform their function even when their ADs are swapped with those of other TFs^21^. This tolerance to diverse AD sequence may be explained by the fact they share the common property of being intrinsically disordered, a feature needed to support phase-separation capacity of TFs^22^. Indeed, the portions of FOXO1, INO80D and NCOA1 involved in PAX3 fusion oncoproteins are predicted to be heavily disordered (Extended Data Figure 5a-c). Remarkably, although transcription factors as a class are heavily disordered (Extended Data Figure 5d), yet PAX fusion partners are particularly disordered (similar to FET family fusions like EWSR1). A related partner MAML3 is almost entirely disordered (Extended Data Figure 5e) and *PAX3-MAML3* fusions occur in biphenotypic sinonasal sarcoma (SNS) but not in RMS^23^. *MAML3* lacks a SE in myogenesis or RMS, and our model would therefore predict the absence of *PAX3-MAML3* translocations in RMS. SNS may arise from a cell of origin whose epigenome has a lineage restricted enhancer at MAML3 which gets recruited in SNS tumorigenesis. Future studies could reveal phase condensate formation as a common capability of all PAX3 fusions found in these tumors.

Many disordered proteins in the genome are not involved in *PAX3* fusions. A parallel criterion for a successfully tumorigenic fusions could be the presence of an active SE in the same lineage step as *PAX3* (such as those SEs proximal to *FOXO1*, *INO80D* and *NCOA1* in the RMS-specific epigenetic state). Super enhancer-containing loci may be enriched in translocations for two reasons. First, active enhancers are transcriptionally active and early replicating^24^, and thus likely more susceptible to double strand breakage^25^. Secondly, among translocations which form, those resulting in overexpression of an oncogene are selected for, and SEs can enable such continued overexpression. The appearance of certain SEs is transient and logically restricted to certain points in development, and thus may be restricting the potential miswiring events that could give rise to an “infinite loop” in CR TF logic. We propose this can explain, at least in part, the selection of translocation partners in FP-RMS tumors, and provides a paradigm likely relevant to other translocation driven cancers.

## Methods

### Circularized chromatin conformation capture (4C-seq)

4C-seq was performed on RH4, RH5 and CTR cells as previously described^9^. Briefly, cells were grown in DMEM at 37 °C, then chemically crosslinked with 1% formaldehyde for 12 minutes. Then, *in-situ* digestion and *in-situ* re-ligation of 3D contacts (keeping nuclei intact) was performed using DpnII as the 4-bp DNA cutter. Following re-ligation, Csp6I was used to reduce template size, followed by relegation to circularize for inverse PCR at viewpoints of interest surrounding *PAX3* and *FOXO1*. 4C samples of RMS cell lines were amplified using bait region primers for PAX3-promoter (F: CAAGGAGTCCTGGTGCCAAA, R: CACTGGCCGGTGAGAAGG). We also studied three CTCF sites on the FOXO1 side with the following primers: FOXO1-CTCF.1 (F: GCTCCCACAGAAGAAGCAGA, R: GGTGGAGACAGAGGCAGTAC), FOXO1-CTCF.2 (F: CACACACAAGCAAGCACAGA, R: AGCCTCATTACCACTTTTGAACA), and FOXO1-CTCF.3 (F: TCAGGAAGGTTCAAACTTACTTTCC, R: CACGCACGCATAAAAGAGCA). Illumina TruSeq ChIP Library Prep Kit was used on purified inverse PCR products, and sequenced (75 bp single end) with an Illumina NextSeq500 (and, are typically multiplexed with ChIP-seq experiments to increase sequence diversity, which is needed because most reads from a single 4C viewpoint are identical duplicated products from self-ligation).

### 4C-seq data analysis

Reads from 4C experiments were first filtered to keep only reads containing the bait primer sequence (from the viewpoint), tolerating 1 mismatched base pair. Then, barcodes and viewpoint sequence were trimmed, followed by mapping to hg19 with bwa. Reads surrounding the viewpoint (within 4 kb) were removed to aid visualization of informative distal contacts. Smoothing was performed by averaging over a sliding window of 5 kb and visualized in IGV.

### 3C confirmation PCR and sequencing

Primers were designed to validate interaction between the outermost CTCF boundaries surrounding *PAX3* and *FOXO1* (Extended Data Figure 1). The left primer overlapped a digestion site detected to form a long-range ligation at high frequency in 4C data, with the sequence: AAGCAGATGGG**GATC**ACGTG (DpnII cutsite in bold). The right primer was designed close to the PAX3 CTCF site, but before any additional cutsites: GCAGCCAGTGAGATAAGACATTA. PCR was performed using Phusion High Fidelity Master Mix (NEB) with 30 pmol of each primer and 70 ng of 3C DNA template (same template from which 4C was prepared, prior to the second Csp6I cut), with 25 cycles (94 °C for 15 seconds, 55 °C for 30 seconds, 70 °C for 30 seconds) with completed with 1 minute at 70 °C. PCR product was purified with PCR QiaQuick kit (Qiagen), then Sanger sequencing was performed at the Center for Cancer Research Sequencing Facility (CCR-SF).

### Exonic imbalance analysis of RMS RNA-seq

RNA-seq read counts were generating using cufflinks at the exon level for each sample (RMS primary tumor data)^7^. Then, for each RMS tumor (n = 97), the exonic balance before and after the known break points were calculated (average FPKM of pre-breakpoint exons/average FPKM of post-breakpoint exons) for all known translocation partners (*PAX3*, *PAX7*, *FOXO1*, *NCOA1*, *INO80D*). Normal exonic bias was corrected for by taking a z-score across all samples (including FN-RMS samples). Plots of exon imbalance where made in R using ggplot2.

### HiC and HiChIP data analysis

HiC data for high-resolution contact frequencies in GM12878 cells^14^ was downloaded and visualized through the Juicebox desktop application^26^. Candidate native loop contacts (purple circles, Figure 1) were annotated manually from visual inspection, and CTCF motif orientation was derived by taking the intersect of CTCF peaks in RH4 cells called with MACS2 (previously published, GEO accession number GSM2214099) and CTCF recognition sequences annotated in the HOMER known motifs collection (http://homer.ucsd.edu/homer/motif/genomeWideMotifScan.html).

H3K27ac HiChIP data analysis of data generated in RH4 cells was performed using the Hi-Pro pipeline^27^. We kept only high-quality paired-end tags (PETs), using the HiC-Pro pipeline to filter out reads lacking the restriction site (DpnII), filter out duplicates, and filter out poorly mapped reads. Valid contact pairs remaining were converted to .hic format using the hicpro2juicebox.sh script (https://github.com/nservant/HiC-Pro/tree/master/bin/utils), to enable compatibility with the Juicebox visualization tool.

### Lentivirus production

A lentivirus plasmid mix was made by combining Lenti-Rev, Lenti-PM2, Lenti-Tat and Lenti-Vsv-G plasmids at a ratio of 1:1:1:2. HEK293T cells were seeded at 50-60% confluency (2 million cells) in a 10 cm dish in 10 mL media (DMEM, 10% FBS) and incubated at 37°C, 5% CO_2_. The next day, 5 μg of transfer plasmid, 10 μg of lentivirus plasmid mix and 36 μL of X-tremeGENE HP DNA transfection reagent (Sigma) were mixed into 1.2 mL serum-free OptiMEM (Gibco), vortexted and incubated for 15 minutes at RT and added to the cells. Media was changed 24 hours after transfection. Viral supernatant was harvested the following three days and filtered through a 0.45 μm syringe filter. Each collection was diluted with 5X PEG-it™ Virus Precipitation Solution (System Biosciences) and stored at 4°C until the final collection. After the final collection, viral supernatants were centrifuged at 1500g for 30 minutes at 4°C. Viral pellets were resuspended in PBS, aliquoted and stored at −80°C. All transductions were conducted with 8 μg/mL polybrene.

### Pooled CRISPR Library Design and Construction

Sequences of enhancer regions (~100 sites) surrounding *PAX3-FOXO1*, CTCF, and SE peaks were derived such that all possible sgRNA guides for *S.pyogenes* Cas9 can be designed within those regions. A total of 1830 sgRNAs were produced. Guides were selected based on having an off-target score of 1.0 denoting the maximum possible score for guides not having any off-target. Oligonucleotides were synthesized as a pool commercially (Twist Biosciences) and then PCR cloned into BsmBI-cut sgRNA expression vector, LRG2.1T, by using gibson assembly. Deep sequencing analysis was performed on illumina platform and verified that all of sgRNA designs were cloned (data not shown).

### Pooled CRISPR-Cas9 screening

The lentiviral sgRNA library was produced as described above. RH4.Cas cells were first transduced with varying concentrations of the pooled lentiviral library to determine the amount needed for a MOI of ~0.3. RH4.Cas cells were then seeded in three 15 cm dishes at 8 million cells per dish. 24 hours later, cells were transduced with the lentiviral library at a low MOI, leading to ~27% positive cells (as determined by FACS analysis of GFP expression) and ensuring over 1000x representation of the library. 48 hours after transduction, virus-containing media was removed and replaced with fresh media. 40 million cells were collected and pelleted at days 2, 5, 7, 10, 13, 17, 20 and 25 post-transduction and 12 million cells were plated at each timepoint for subsequent cell sampling. Coverage at cell level was kept above 1000x throughout the screen.

Sequencing libraries were constructed essentially as previously described (Shi et al., 2015). Genomic DNA (gDNA) from cell pellets was extracted using the AllPrep kit (Qiagen). 32 parallel PCR reactions were performed to amplify sgRNA sequences from gDNA harvested at each timepoint. The total volume of each PCR reaction was 50 μL containing 400 ng of gDNA template, 0.2 μM forward (TCTTGTGGAAAGGACGAAACACCG) and reverse (TCTACTATTCTTTCCCCTGCACTGT) primers, and 25 μL Platinum PCR SuperMix (Thermo Fisher). PCR cycles were: 1x (98°C, 2min) 35x (98°C, 8s, 67°C, 12s 72°C, 10s), 1x (72°C, 5min). Parallel PCR products were pooled together and purified with the QiaQuick kit (Qiagen). After production of sgRNA amplicons from gDNAs by PCR, ends of fragments were repaired using T4 DNA polymerase (NEB), DNA polymerase I large fragment (Klenow) (NEB), and T4 polynucleotide kinase (NEB), followed by addition of 3’ A-overhang with Klenow (3’-5’-exo-) (NEB). Amplicons from each sample were ligated with a unique barcode adaptor (pool) and purified using AMPure magnetic beads (Bechman Coulter). Samples were deep sequenced on Illumina platform. Read counts for each sample were ascertained by mapping raw reads to the sgRNA library sequences. Read counts of each sample were normalized to facilitate data analysis.

### Construction of individual sgRNA expression plasmids

sgRNA sequences were designed using the CRISPR design tool from MIT (crispr.mit.edu). Pairs of DNA oligonucleotides encoding the protospacer sequences were annealed together to create double-stranded DNA fragments with 4-bp overhangs. These fragments were ligated into BsmBI digested Shuttle_sg_RFP657 plasmid. Plasmid constructs were confirmed via sanger sequencing using the LKO1_5 primer (GACTATCATATGCTTACCGT) on the U6 promoter.

## Data Access

Newly generated ChIP-seq data from primary RMS tumors has been made publicly available through the Gene Expression Omnibus (https://www.ncbi.nlm.nih.gov/geo/). The GEO accession number is GSE136799. Code is available at https://github.com/GryderArt.

## Acknowledgments

We thank Michael Kuehl and Katherine Masih for thoughtful discussions of the manuscript. We thank the CCR Genomics Core at the National Cancer Institute, NIH, Bethesda, MD for Sanger sequencing. This work was supported by the NCI, NIH. Silvia Pomella is a recipient of a Fondazione Veronesi fellowship.

The content of this publication does not necessarily reflect the views or policies of the Department of Health and Human Services, nor does mention of trade names, commercial products, or organizations imply endorsement by the U.S. Government.

