## Supplementary Figures 1-5 for "Miswired enhancer logic drives translocation positive rhabdomyosarcoma"

**Supplemental Figures and Legends**

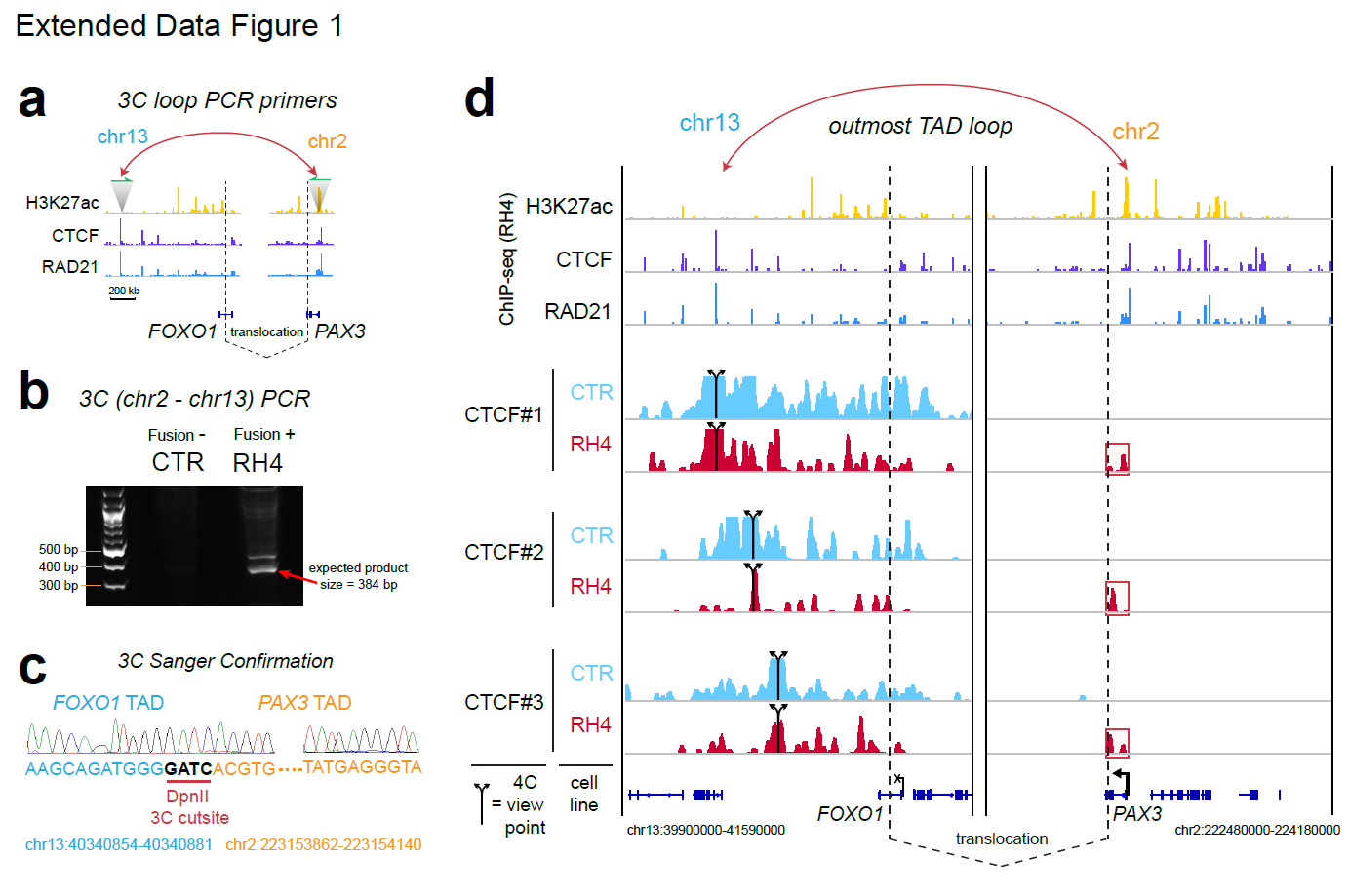

**Extended Data Figure 1**

**a.** Diagram of 3C confirmation PCR primer placement

**b.** 3C confirmation PCR in fusion negative (CTR) and fusion positive (RH4) cells.

**c.** Sanger validation of 3C PCR product.

**d.** 4C-seq of CTCF sites: 3 viewpoint sites distal to *FOXO1* on chromosome 13, which show interaction with chromatin proximal to the *PAX3* promoter and CTCF anchor.

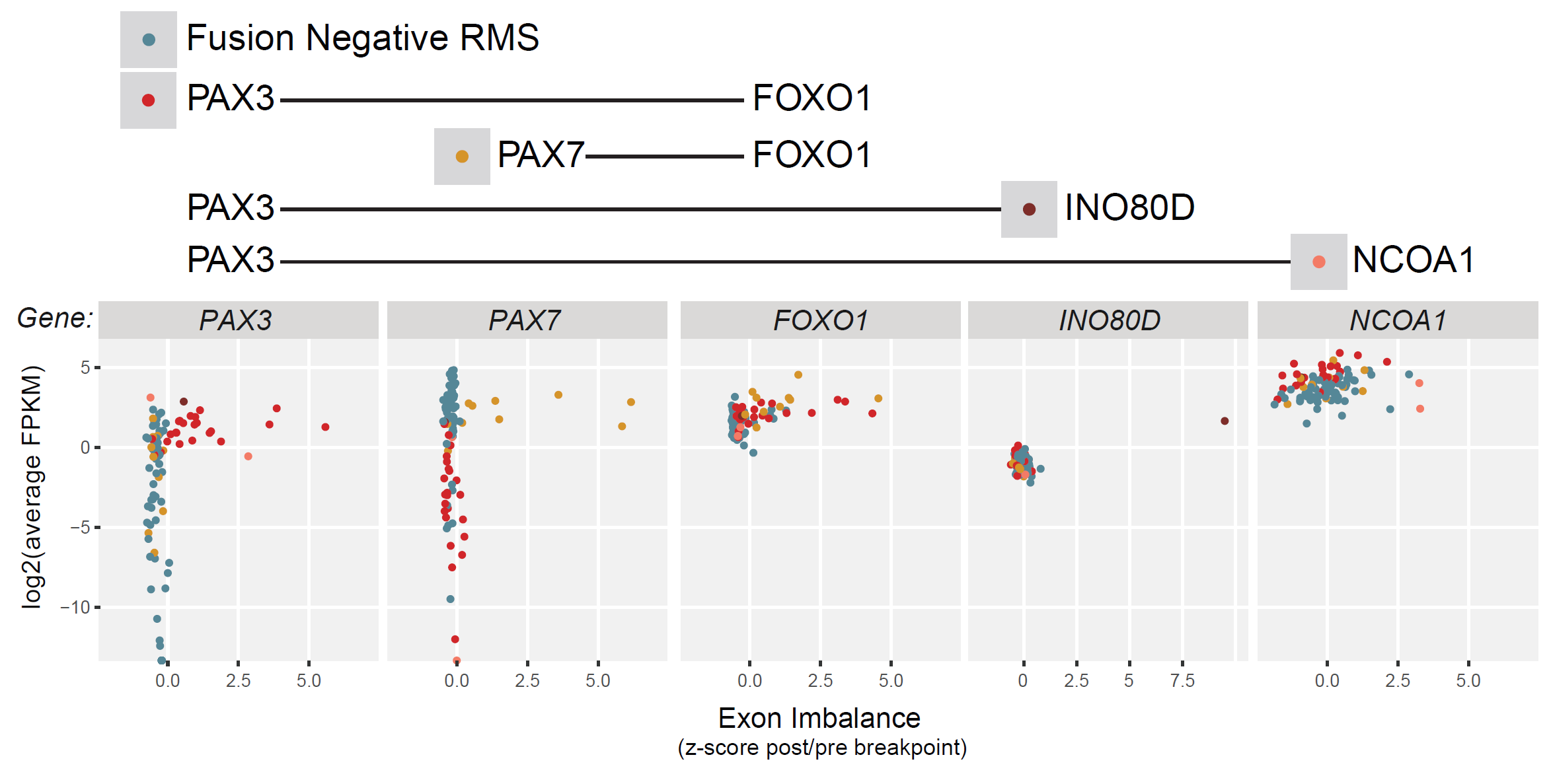

**Extended Data Figure 2**

Exon imbalance for PAX fusion partner genes, plotted against RNA expression levels.

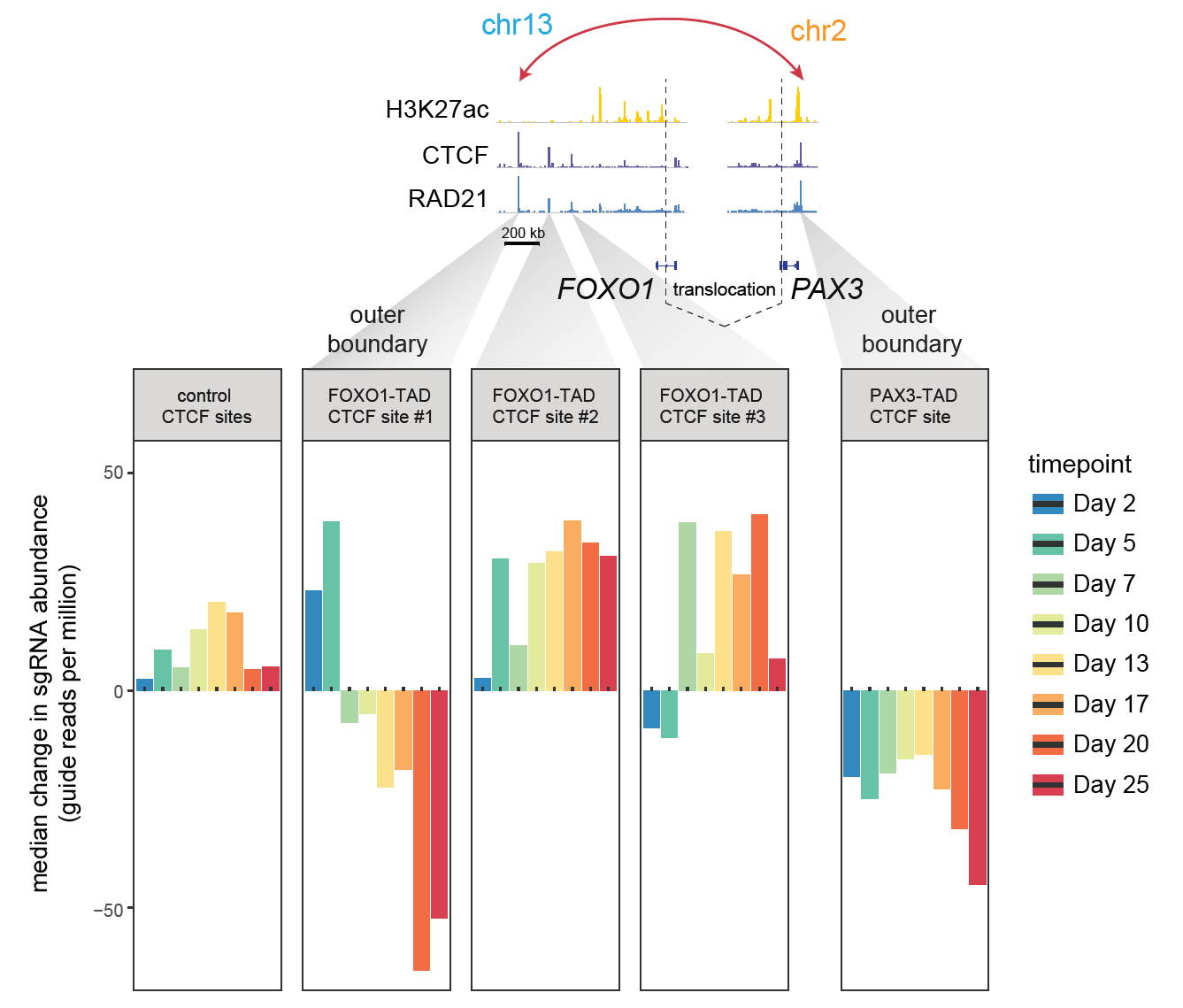

**Extended Data Figure 3**

Pooled sgRNA CRISPR interrogation of CTCF sites surrounding *PAX3* and *FOXO1* reveals importance of the two most distal sites, but not two intervening CTCF bound locations distal to *FOXO1*.

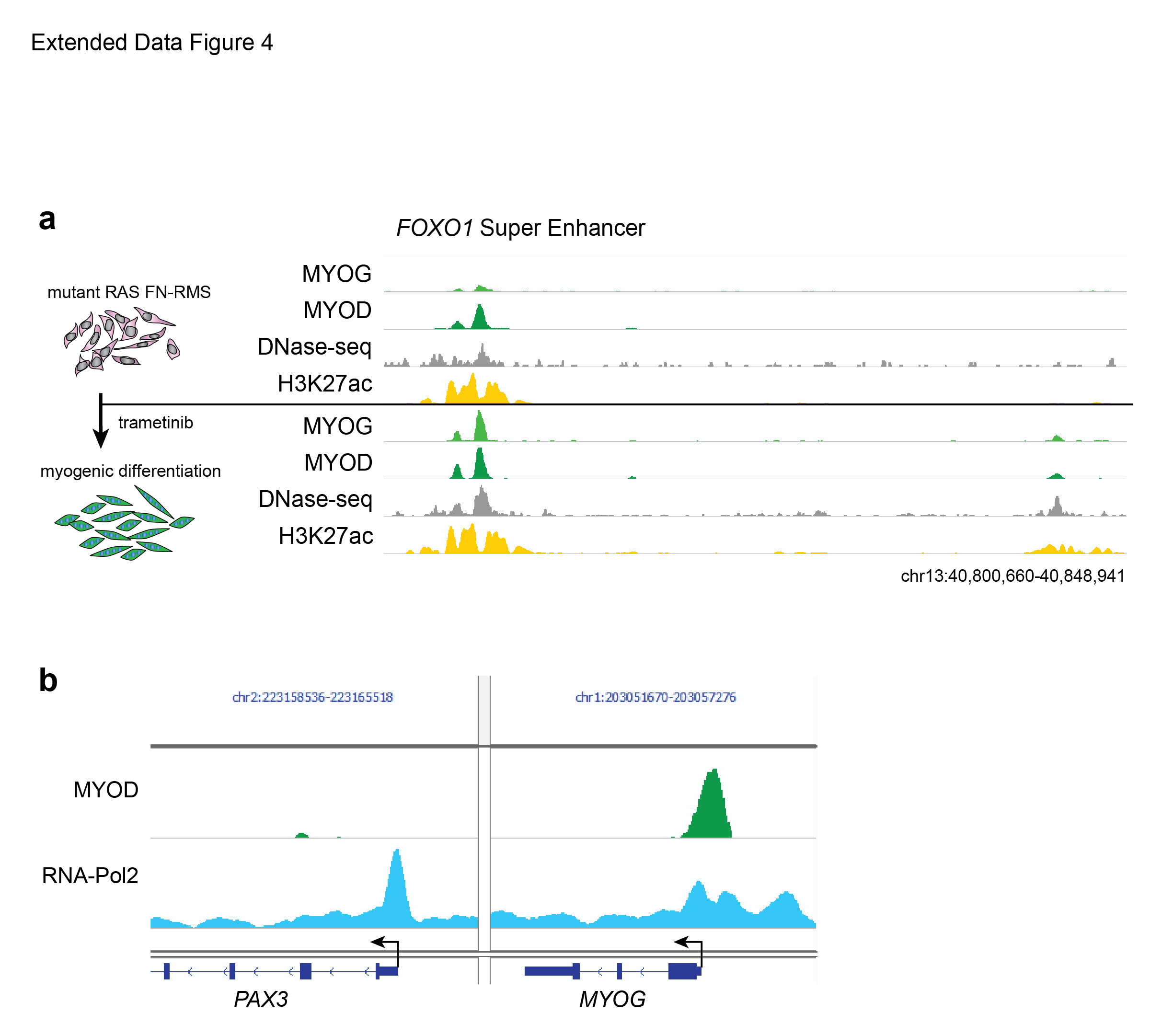

**Extended Data Figure 4.**

a. MYOG invades the *FOXO1* super enhancer during trametinib-induced myogenesis in SMS-CTR cells, a PAX-fusion lacking cell line driven by mutant HRAS (Q61K).

b. MYOD ChIP-seq alongside RNA Pol2 ChIP-seq at the *PAX3* and *MYOG* promoters in RH4 cells (*PAX3-FOXO1* translocated FP-RMS cancer cells)

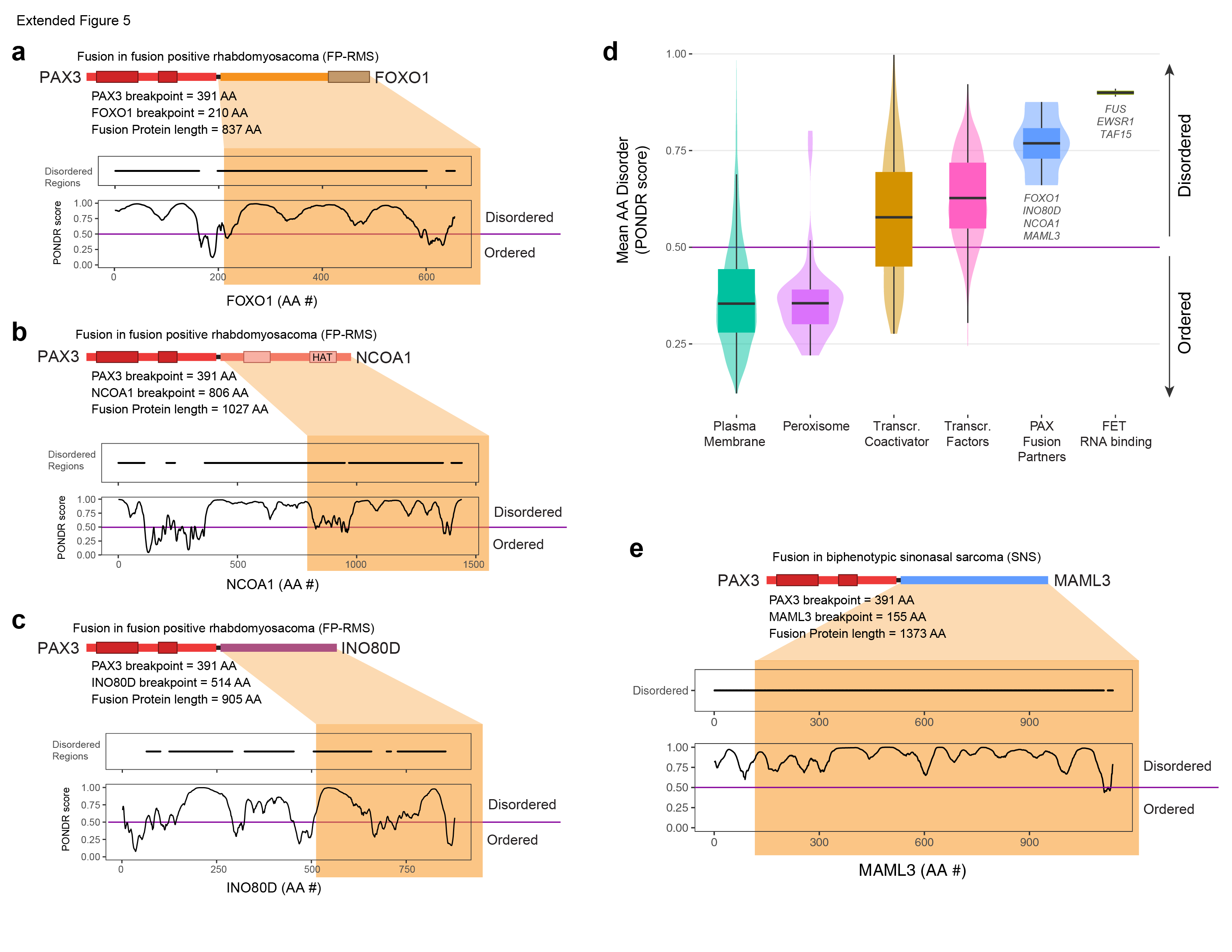

**Extended Data Figure 5.**

**a.** PAX3-FOXO1 fusion gene, with PONDR (Predictor of Natural Disordered Regions) score (<http://www.pondr.com/>) and disordered regions mapped for FOXO1 amino acids.

**b.** PAX3-NCOA1 fusion gene, with PONDR score for NCOA1.

**c.** PAX3-INO80D fusion gene, with PONDR score for INO80D

**d.** Distribution of average disorder PONR scores among protein families, including transcriptional coactivators and transcription factors, alongside PAX3 fusions and FET family fusion partners.

**e**. PAX3-MAML3 fusion gene, with PONDR score for MAML3.
